# Traveling brain waves support flexible storage of human working memory

**DOI:** 10.64898/2026.10.01.755588

**Authors:** Yifan Zeng, Yannik Hilla, Charline Peylo, Paul Taylor, Lukas Imbach, Johannes Sarnthein, Andrea Alamia, Paul Sauseng

## Abstract

Human working memory enables us to temporarily store and flexibly utilize information. It is thought that there are two distinct neural mechanisms by which information is retained in working memory: persistent firing of working memory-relevant neurons^1^, or short-term synaptic plasticity in the absence of ongoing neuronal firing^2^. However, both these mechanisms can lead to strong neural interference when working memory contents are updated with novel information. As an alternative, here we propose that information can be flexibly held in working memory in the form of traveling waves - spatiotemporal patterns of neural activity that propagate across brain tissue. In an artificial neural network, we simulated wave propagation by passing transient neural firing patterns representative of single-item information across different network layers. In comparison to sustained neural firing, we demonstrate that these transient traveling waves result in minimal synaptic plasticity, thereby facilitating working memory updating without significant neural interference. We then identified such traveling waves in intracranial recordings from the human temporal lobe. Traveling waves precisely when information was being encoded and persisted uniformly throughout maintenance and recall, aligning with the activation pattern seen in the artificial neural network. In trials where participants committed memory errors, no such continuous traveling wave activity in the temporal lobe was observed. Our data from artificial neural networks and human intracranial brain recordings converge to indicate that neural traveling waves support information processing in working memory while minimizing interference during updating. This here reported mechanism, together with the classical working memory models, serves as a mutually complementary, multi-mechanistic framework for working memory storage.

## Main

Humans are highly efficient at transiently storing sensory information in memory, and flexibly updating it to guide their behavior. Together, these processes are known as working memory^3^– a cognitive function essential in our daily lives. To date, there are two major theories on how working memory is implemented in the brain: (i) sustained firing of neural assemblies coding for information in working memory^4^ and (ii) activity-silent coding based on short-term and very transient synaptic plasticity^2^. Whereas both of these theories can very convincingly explain how information is transiently stored in working memory, they cannot fully account for how content is flexibly updated within working memory: Both sustained neural firing that causes synaptic weight changes^5^ and activity-silent synaptic plasticity can lead to strong interference between outdated and novel information in working memory^6^. Here, we suggest a third, additional neural mechanism by which working memory is implemented in the human brain– a mechanism that minimizes updating-related interference based on activation patterns traveling across brain structures.

If you throw a stone into a pond, you will see waves propagating away from the location of impact. By the time these waves reach your position at the bank, they reflect an event that has happened some time in the past (namely the stone hitting the water surface), and therefore may be considered as a memory trace of an earlier event^7^. Such traveling waves, i.e., mesoscopic neural activity patterns that propagate spatially across the brain, have been observed in the rodent^8,9^, non-human primate, and human brain^10–13^, but their functional role remains unclear. Previous work linked traveling waves to sensory processing^14,15^ and perceptual detection^16^. Traveling waves were also found to be associated with working memory encoding^17,18^, maintenance^17,19^, retrieval^11,17,18,20^, and filtering^21^, as well as sleep-dependent consolidation^22,23^. Importantly, one recent study directly proposed traveling waves as a potential mechanism for shielding working memory from distracting information^19^. But could such traveling waves also be involved in the *transient and flexible buffering* of information per se?

Persistent activation of a defined neural population may result in substantial changes in synaptic weights^24,25^. Indeed, synaptic plasticity is considered a way in which information can be transiently stored in working memory^2,26^. However, such hard-wired changes in synaptic weights are difficult to reconcile with the flexibility of working memory, when contents must be rapidly updated and replaced as task demands evolve^27,28^. By continuously shifting the location of neural activity and thereby reducing repeated stimulation of individual neurons and synapses, traveling waves provide the possibility of a mechanism for information storage that is less dependent on synaptic plasticity. This is why we hypothesize that in a flexible working memory system, neural representations of retained items would be passed on between neurons like a traveling activation pattern, similar to a wave in a pond; i.e., a chain of neurons would be used as a notepad for buffering information. As single neurons are activated only transiently before passing information on to the next neuron in line, traveling waves avoid the hardwiring of specific information into synaptic connectivity, making this mechanism more robust against interference, particularly during working memory updating.

We tested this idea by setting up a simple computational network model in which we compared how well information of multiple items held in working memory was stored and updated by (i) stationary, persistent activity of network nodes versus (ii) neural activity being passed on from network node to node in a traveling activation pattern. Then, we hypothesized that the node-level activity observed in the simulations is sufficiently consistent across the neuronal population to give rise to similar traveling neural activity at the level of the local field potential. We thus analyzed intracranial electroencephalography (iEEG) recordings from patients with implanted electrodes in the temporal lobe while they were doing a working memory task. Specifically, we examined traveling wave patterns to test whether similar propagation of neural activation, as predicted by our simulations, can be found in the human brain.

### Simulation of traveling neural activity as a substrate for working memory storage

For illustration purposes, we built a computational network model consisting of 150 nodes organized into a unidirectional ring (Fig. 1A). The network consisted of 15 layers of 10 nodes each, where the nodes of each layer (y) were sequentially and fully connected to those of the subsequent layer (y+1). The last layer was fully connected to the first layer, resulting in a unidirectional ring (Fig. 1 shows a simplified illustration with only four layers and three nodes each). The connections between neighboring layers followed a Hebbian-like behavior and were strengthened when presynaptic and postsynaptic nodes fired together (see Methods for details).

**Figure 1.**
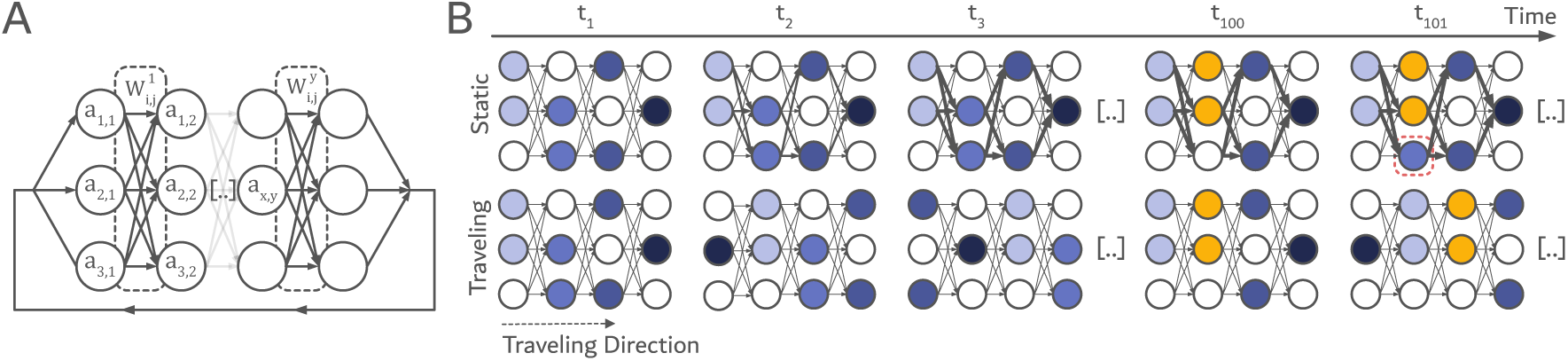
The computational network model of static and traveling working memory storage. (A) Architecture of the artificial neural network. Adjacent layers were fully connected, and synaptic weights were updated according to a Hebbian learning rule. The nodes in the final layer were connected back to the ones in the first layer, forming a ring connection. (B) Conceptual illustration of the static and traveling memory models. In the static model, stored neural representations remain at the same network locations over time. In the traveling model, neural representations propagate to the next layer at each time step. We hypothesize that the static model develops strong synaptic connections through repeated activation, leading to increased interference when stored information is updated (at t = 100). In contrast, the traveling model relies less on synaptic plasticity because representations continuously move through the network, thereby reducing interference and enabling more flexible memory storage.

With this network, we tested two different coding scenarios: (i) static activation coding (Fig. 1B, upper panel), and (ii) traveling activation coding (Fig. 1B, lower panel). In both models, single items were always represented by the activation code within different layers. In the static activation scenario, activation stayed at the initially stimulated nodes in the network at any time point. In the traveling activation coding scenario, the initial activation pattern propagated to the next downstream layer at each time step (activation in the last layer circled back to the first layer due to the ring network structure). This means that, at each time step, an activation code representing a particular item stored in memory was passed on from one layer to the next. Importantly, we hypothesized that sustained activation of the same node pairs in the static condition promoted Hebbian strengthening of synaptic connectivity (indicated by strengthened connections between certain nodes in Fig. 1B, upper panel), whereas the continuously propagating activity pattern of traveling waves distributed activation across the network, thereby limiting synaptic changes between specific node pairs. To simulate working memory updating, the activation states of a subset of nodes were updated after 100 time steps, representing the replacement of some memory items with new ones. As compared to the traveling model, greater synaptic change was expected in the static model. Consequently, previously established synaptic weights were expected to exert stronger interference during memory updating.

To test this hypothesis, we examined memory maintenance and updating in the static and traveling-wave models while varying the Hebbian learning rate (i.e., the parameter determining how strongly synaptic weights change in response to correlated neural activity; δ = 0.05 or 0.1). As an indicator of how well the two models retained the correct memory information, the Hamming distance between network activation patterns at any given time step and the ground truth was computed (Fig. 2A, light and dark red curves). The smaller the Hamming distance, the better the model’s performance in representing the exact neural code for memory items. As shown in Fig. 2A, over time, the Hamming distance increased in the static model but barely in the traveling model. While the deviation from the ground truth in the first 100 time steps was likely due to the random weight initialization (see Methods), these results indicated that the traveling model is superior in retaining the initial information against noise in the weights.

**Figure 2.**
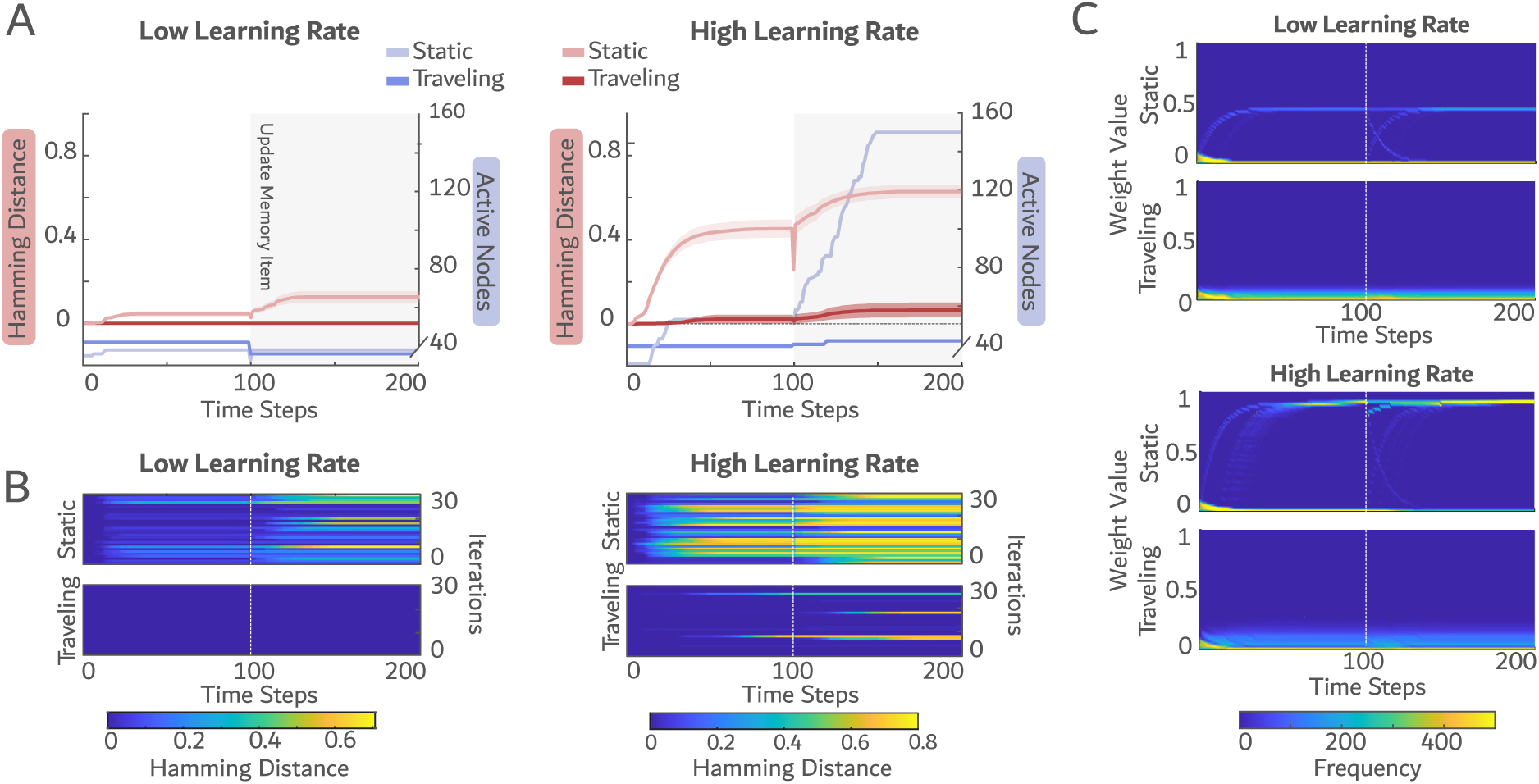
The traveling model outperformed the static model. (A) Temporal dynamics of the two models’ Hamming distances from the ground truth and the total number of active nodes in the network for two learning rates (δ = 0.05 and 0.1). Shaded regions indicate the standard error across iterations. (B) Hamming distances in all 30 iterations for each model and learning rate condition. (C) Temporal evolution of the synaptic weight distribution.

At time step 100, memory content was updated, i.e., coding of some memory items was replaced by new ones. This resulted in a striking increase in deviation from the ground truth in the static model, whereas the traveling model showed only minimal disturbance, even under high learning rates. To assess the consistency of model performance, we repeated the simulation for 30 iterations, each with different initializations. While the traveling model showed notable deviation from the ground truth in only a few iterations at a high learning rate, the static model exhibited substantial Hamming distances from the ground truth across most iterations (Fig. 2B).

Similar effects as for Hamming distance were obtained for the total number of activated nodes in the network. As can be seen in the blue curves of Fig. 2A, both the traveling and the static models produced a low and constant number of activations over the 200 time steps when the learning rate was low. However, a higher Hebbian learning rate led to a high perturbation exclusively in the static model, where the activation increased dramatically following memory update at time step 100. In contrast, the traveling model remained largely unaffected.

As expected, the distribution of synaptic weights indicated that in the traveling model there was hardly any synaptic change, neither under low nor under high learning rate (Fig. 2C). In contrast, synaptic weights in the static model progressively saturated, with many approaching their theoretical maximum. This finding underpins the assumption that, in the traveling model, synaptic plasticity during memory maintenance and updating is largely prevented. Reduced synaptic plasticity mitigates interference during memory updating; theoretically this may particularly be the case for neural structures that exhibit high learning rates, such as the medial temporal cortex^29^. This motivated the next step of the study, where we analyzed local field potential (LFP) data recorded invasively from the medial temporal cortex of patients with epilepsy while they were performing a working memory task^30^.

### Traveling theta and gamma waves in the temporal cortex during a working memory task

A publicly available LFP dataset^30^ of 15 participants with focal epilepsy (age = 31.8 ± 12.9 years; 8 females and 7 males; 3 of 15 participants were excluded from the current analysis for not having enough electrodes, leaving n = 12; see Methods) was re-analyzed. LFPs were obtained via implanted depth electrodes in the lateral and medial temporal cortex (Fig. 3B) while participants performed a Sternberg task. In each trial, four, six, or eight consonants were presented for 2000 ms; participants were instructed to maintain the consonants in memory for an additional 3000 ms and subsequently indicate whether a probe consonant had been part of the memorized letter string (Fig. 3A). Participants performed the task well, achieving median correct-response rates of 98.1% (IQR: 96.9–99.6%) for set size 4, 92.0% (IQR: 89.4–94.1%) r set size 6, and 85.5% (IQR: 79.3–90.5%) for set size 8. Median reaction time increased with memory load (48.3 ms/item), from 1,087 ms (IQR: 971–1,194 ms) for set size 4 to 1,141 ms (IQR: 971–1,194 ms) for set size 4 to 1,141 ms (IQR: 1,089–1,297 ms) for set size 6 and 1,244 ms (IQR: 1,167–1,321 ms) for set size 8, consistent with previous studies^31^.

**Figure 3.**
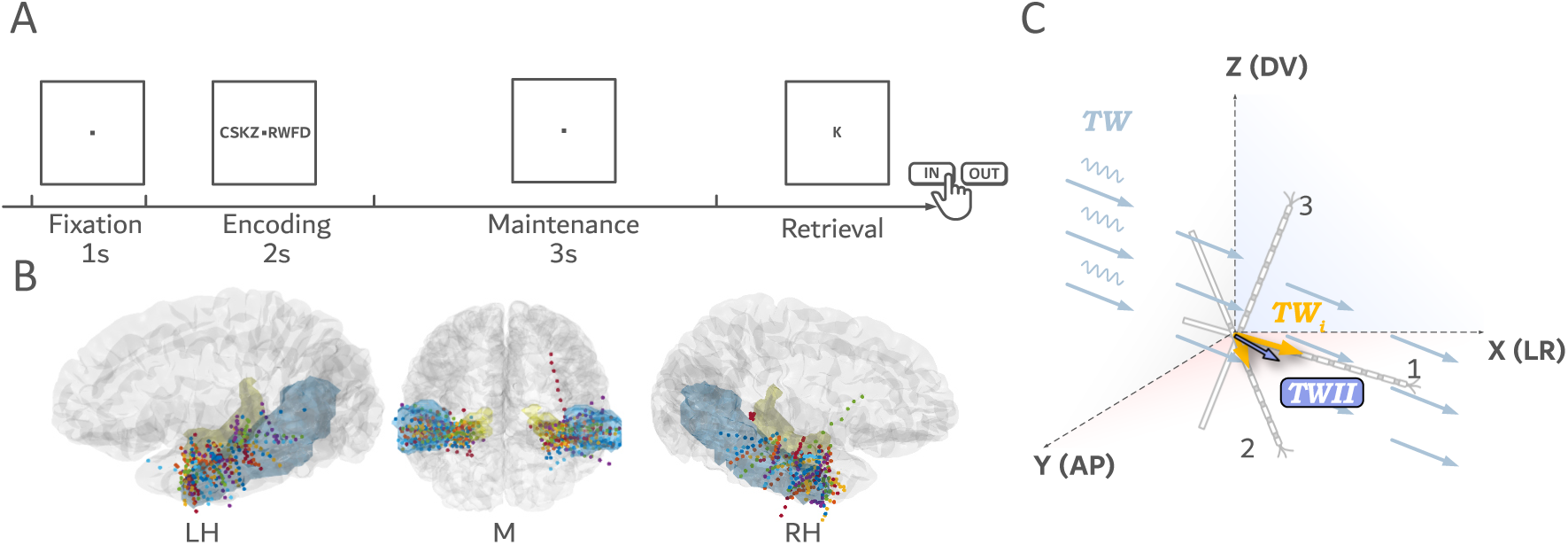
The dataset from Dimakopoulos et al. (2022)^30^and traveling wave computation. (A) The verbal Sternberg task. (B) Positions of contacts on the implanted depth electrodes in middle temporal gyrus (blue) and hippocampus (yellow), collapsed across participants. (C) Computation of TWII. We first computed traveling wave intensities for each electrode, and then aggregated them according to their orientations in space. See “Method: Traveling Wave Intensity Index” for more details. LH, left hemisphere; M, middle view; RH, right hemisphere; TW, true traveling waves in the region; TW_i_, traveling wave intensity acquired by an individual electrode; TWII, traveling wave intensity index in the region.

In this dataset, we aimed to test whether – as suggested by the above-described artificial neural network – traveling neural activation patterns were involved in encoding, maintenance, and retrieval of information in working memory in brain areas known for showing high synaptic plasticity and a high synaptic learning rate. To simultaneously quantify direction and intensity of traveling waves (i.e., the consistency of the phasic gradient across contacts), we introduced the Traveling Waves Intensity Index (TWII). TWII is a three-dimensional vector: its orientation indicates the estimated traveling wave direction, and its magnitude indicates the consistency of the propagation. A TWII with a magnitude of zero indicates the absence of consistent traveling waves, whereas larger magnitudes correspond to more consistent traveling waves propagating along the orientation of the TWII. Fig. 3C illustrates the process for computing the TWII in the temporal lobe. We first quantified the intensity of traveling waves at each temporal frequency along each individual multi-contact electrode (noted in Fig. 3C as electrode 1, 2, and 3). This was achieved using a method based on the 2D fast Fourier transform ^13,21^, yielding one vector for each time-frequency point and each electrode that represented the net direction and intensity of traveling waves along the electrode (Fig. 3C, yellow arrows “TW_i_”). We then aggregated these vectors across electrodes within the same hemisphere by performing a vectorial mean, and this gave a TWII that represented the regional traveling waves for a given time-frequency point (Fig. 3C, the purple arrow). We applied a weighted 3D Rayleigh test (developed from the 2D weighted Rayleigh test^32^) to identify time-frequency clusters of traveling-wave patterns across all participants. Thereby, a significant cluster-level statistic, R*, indicates that traveling waves within a cluster are consistently expressed across participants beyond chance level. This way, in correctly responded trials, it was possible to obtain multiple time-frequency clusters in both hemispheres (Fig. 4A, upper panel): In the left hemisphere, one continuous cluster in the theta frequency range between 4 and 10 Hz, as well as several clusters at gamma frequency between 32 and 56 Hz were identified. Strikingly, traveling waves at both frequency bands had a sharp onset at presentation of the memory set (i.e., the stimulus string), and were then sustained throughout the encoding stage, the maintenance stage, and memory retrieval. After retrieval of the information from working memory, the consistent traveling waves vanished (particularly abruptly at theta frequency; Fig. 4A, upper-left panel). Importantly, throughout encoding, maintenance, and retrieval, traveling theta and gamma waves propagated consistently in the same direction (Fig. 4B, left panel), from the lateral temporal cortex towards the medial temporal lobe. This continuous and uniform traveling of theta and gamma waves during all working memory processes are reminiscent of traveling patterns in the above-described artificial neuronal model, where neural activation patterns were passed on from one layer of neural nodes to the next one.

**Figure 4.**
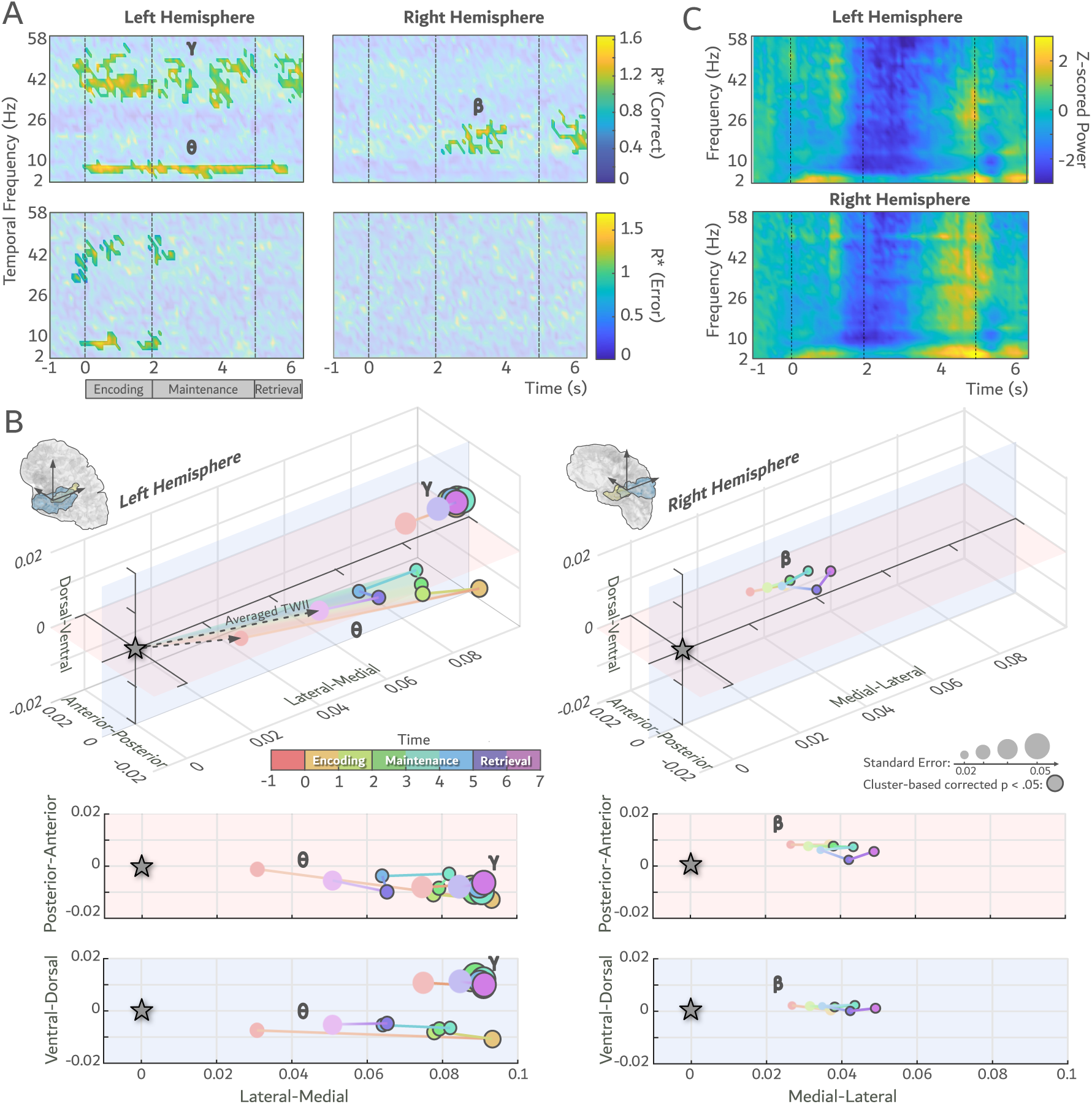
Theta, beta, and gamma waves traveling in the temporal cortex during working memory. (A) The temporal dynamics of the traveling waves separately for trials with correct (upper panel) and error responses (lower panel). The highlighted regions show time-frequency clusters that reveal consistent traveling wave patterns across participants. All those traveling activations were disrupted or disappeared in error trials. (B) The direction of the traveling waves in three frequency bands as indicated in (A). The trajectories of the head of TWIIs are depicted, with their tail always centered on (0,0,0). (C) The temporal dynamics of Z-scored wave power (averaged FFT power across all contacts). γ, the gamma traveling waves in the left hemisphere; θ, the theta traveling waves in the left hemisphere; β, the beta traveling waves in the right hemisphere;

In the right hemisphere, two clusters of consistent traveling waves in the beta frequency range between 14 and 30 Hz appeared (Fig. 4A, upper-right panel). The right hemispheric beta frequency cluster reached significance only during the first half of the maintenance stage and then during retrieval of information when the probe item was presented. Moreover, right hemispheric beta waves propagated from more medial towards lateral temporal cortex (Fig. 4B, right panel). We here focus on the TWII observed in the left hemisphere, as we found a left hemispheric dominance for this language-related task where all participants were right-handed^30,33^.

The estimated average traveling speed of theta, beta, and gamma waves in the temporal cortex varied from 0.18 to 1.09 m/s (See Supplementary Analysis; Extended Data Fig. 2B) and is therefore in a range comparable to previously observed mesoscopic traveling waves^34^. Theta and beta activations throughout the trial had a power magnitude higher than the expected 1/f distribution (see Extended Data Fig. 2A). The temporal dynamics of the Z-transformed power for all frequencies (Fig. 4C) were independent of the pattern observed for traveling waves (Fig. 4A). This indicates that the observed traveling wave dynamics were not merely driven by any amplitude fluctuations throughout the task.

### Continuous theta and gamma traveling waves in the left temporal cortex are linked to successful working memory storage

If - as indicated by the suggested artificial neural network model and intracranial recordings from human temporal cortex - traveling theta and gamma waves are a signature of successful encoding, maintenance, and retrieval of information in working memory, one would expect that these signatures are attenuated in unsuccessful trials of a working memory task. To test this hypothesis, TWII was calculated separately for incorrectly answered trials. As depicted in the lower-left panel of Fig. 4A, initially during the encoding stage and around the transition from stimulus offset to maintenance onset, significant theta- and gamma-band traveling waves were observed. These waves occurred at frequencies comparable to those observed in correct trials and propagated in similar directions across participants. However, in stark contrast to correct trials, they were not sustained during error trials; throughout the maintenance and retrieval stages, no consistent traveling waves were detectable. Due to unbalanced trial numbers between correct and error trials, we tested a direct comparison between trial types using a Bayesian Analysis, fitting a model with time and response type (correct vs. error) as predictors of significance in Fig. 4A. The results showed that an interaction between response type and time explains a greater portion of the variability in traveling wave significance than the model with only main effects (theta band [8Hz] BF_10_ > 100, gamma band [40Hz] BF_10_ = 41.95). This indicates that, in line with the above-outlined mechanism for how information could be transiently stored in working memory, errors tend to occur when theta- and gamma-band waves fail to propagate continuously from lateral to medial sites throughout the working memory task.

The artificial neural network model presented above suggests that storing information by passing on neural codes as continuous traveling activation patterns can make working memory more robust to perturbations (e.g., updating of memory content). A similar effect was observed in the data recorded from the human temporal lobe. Only with continuous traveling waves propagating in the same direction throughout the entire trial (i.e., encoding, maintenance, and retrieval stages), information was successfully stored in working memory, and correct responses were given by participants. Discontinuous traveling waves, lasting only for brief periods around encoding and early maintenance, were associated with incorrect responses. This highlights the relevance of traveling waves for working memory processes, in line with recent work^17–21^. Depending on the spatial distribution of these traveling waves, their exact function for working memory might differ. They might enable sequential “scanning” through visual space during working memory processing ^20^, coordinate between brain regions during working memory-guided behavior ^35^ and protect memory content from distracting information ^21^. Here, however, we demonstrate that traveling waves can constitute a neural mechanism through which information is maintained in working memory *per se*^19^.

### The directions of traveling waves align with the information flow during working memory processes

In the left hemisphere, we observed waves propagating from the middle temporal gyrus to the hippocampus in the same hemisphere consistently throughout the entire working memory process. In agreement with the original research of the online dataset, the direction of traveling waves aligns with information flow during verbal memory encoding, i.e., first the semantic processing in the primary auditory cortex and further passed on to the hippocampus^30^. For continuous traveling waves propagating in the left hemisphere to carry working memory information, neural activity cannot simply follow a one-way trajectory from the lateral temporal cortex to the medial temporal cortex, with the neural code being lost upon reaching the medial temporal lobe. Instead, recurrent direct fiber connections returning from the medial temporal cortex back to the lateral temporal cortex are required to make these structures comparable to our artificial neural ring model. In other words, via direct connections, neural information is required to be fed back from the medial temporal cortex into lateral temporal areas to trigger the next traveling wave propagating on the lateral to medial axis. From our data, we cannot conclude whether this truly is the case. However, the acoustic radiation and the inferior longitudinal fasciculus seem plausible candidate fiber bundles for returning activity from the medial temporal lobe to the lateral cortex^36^.

### Theta and gamma waves propagate concurrently and simultaneously in the same direction

It is noteworthy that the working memory-relevant traveling waves throughout the entire memory trials are in the theta and the gamma frequency range. These two frequencies, particularly in interaction with each other, have been frequently associated with working memory processes^37^. Specifically, theta-gamma coupling has been proposed as the physiological basis of a multi-item working memory buffer, where different memories are thought to be stored in different gamma cycles nested into different phases of a repeating theta cycle ^38,39^. Cortical and hippocampal theta-gamma coupling has been observed across a variety of working memory tasks with systematic variations across different memory loads, fidelities, learning stages and performances^40–48^.

## Conclusion

We suggest that information can be transiently stored in working memory via traveling neural activity waves at theta and gamma frequencies. This allows minimal dependency on neural plasticity, hence more flexible storage and less interference when updating new information. The newly suggested traveling wave mechanism, together with classical working memory models that rely on synaptic changes or sustained stationary neural activity, serves a mutually complementary, multi-mechanistic framework for working memory storage.

## Methods

### Neural Network Simulation

#### Architecture

The network consists of 150 nodes, organized into 15 layers of 10 nodes each. Each node can be in an active or inactive state, and these are randomly initialized in the first time step with a value a_x,y_ = {0, 1}, such that a given percentage of nodes are in the active state. A node can change states following the dynamics described below. The nodes of layer *y* are sequentially and fully connected to those of layer *y+1*. The last layer is also fully connected to the first layer in a unidirectional ring network (Fig. 1A). The connections are defined by a connectivity matrix 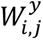 where y determines the input layer, and i,j are the indices of the nodes at the layer y and y+1.

#### Network dynamics

The network is initialized to have 25% of the nodes in the active state at the first timestep. Over timesteps, each state a_x,y_ of each node x at the layer y is updated according to the following formula:

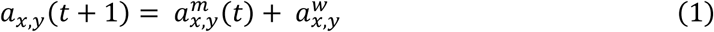

The term 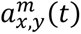 represents the activity that maintains the relevant information in each node. We consider a ‘traveling’ and a ‘static’ condition, defined as:

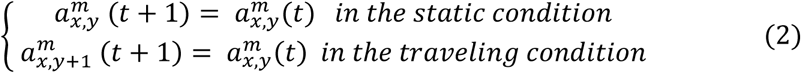

In the traveling condition, the 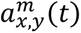 term propagates from layer y to layer y+1, but not in the ‘static’ condition. The second term 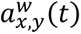 determines the effect of the weights, and it is defined as the product between the activity of the layer y-1 and the weights 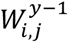 between layer y-1 and layer y:

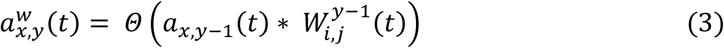

Where *Θ*(*x*) is defined as

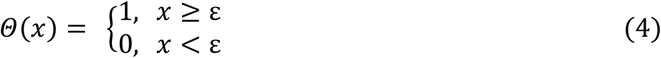

Note that a node can change its state over timesteps as schematically shown in Fig. 1B. Importantly, a node can switch from inactive to active due to the effect of the connectivity matrix, if 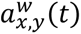 is larger than the threshold ε, which was set to 0.75.

#### Weights update

The connectivity matrix 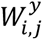 was updated at every timestep following the equation:

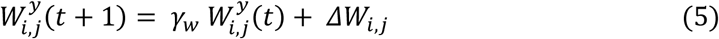

Where γ*_w_* is a decay term we set to 0.9, and *ΔW_i,j_* followed a Hebbian-like behavior defined as:

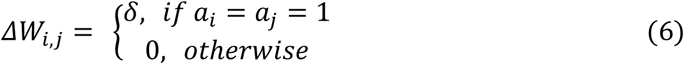

Where δ was set to 0.05 or 0.1 to investigate the behavior of the network with different learning strengths.

#### Procedure

At first, we initialized the network such that 25% of the nodes were in the active state. The weights 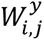 were initialized randomly from a Uniform distribution *U(0,0.1).* We run the network for 200 timesteps. Importantly, after 100 timesteps, we modified the activity of eight layers, setting the states to new, random values, thus mimicking the updating of the encoding of some items with new ones. We quantified the encoding of activity in the layer over timesteps by measuring the Hamming distances between the expected activity (i.e., if there were no interference from synaptic connections) and the network activity. We also quantified the changes in the synaptic weights over time steps in the two conditions by assessing their distribution at each timestep. We repeated the procedure thirty times with random initializations.

### Human Intracranial Dataset

We reanalyzed a publicly available dataset from Dimakopoulos et al. (2022)^30^ (accessible at https://openneuro.org/datasets/ds004752/versions/1.0.1), which includes recordings from 15 participants who underwent depth electrode implantation. Local field potentials were recorded while participants performed a modified Sternberg task. Below, we briefly summarize the methods used in Dimakopoulos et al. (2022)^30^ and describe how it was utilized in our reanalysis. For comprehensive details of the dataset, please refer to the original publication.

#### Participants

The 15 participants of Dimakopoulos et al. (2022)^30^ were patients with focal epilepsy (age = 31.8 ± 12.9, 8 females and 7 males). The intracranial electrodes were implanted in all the participants to investigate a potential surgical treatment for refractory epilepsy. All the patients were under medication. Data from three participants were excluded from the current reanalysis due to insufficient depth electrodes (see “Traveling Waves Intensity Index”). This leads to 12 participants in the present study (age = 32.6 ± 12.9, 6 females and 6 males). The patients gave written consent for the original study (ethical number: PB 2016–02055); the dataset is publicly available and no additional ethical approval was acquired for the current reanalysis.

#### Experimental Task

Dimakopoulos et al. ^30^ used a modified verbal Sternberg task (Fig. 3A). Each trial starts with a fixation of 1s, followed by the presentation of a list of letters at the center of the screen. Four, six, or eight consonants were presented in the middle as memory items. The set size of memory items was randomized across trials. Eight letters were always presented to ensure a similar amount of visual stimulation for all set sizes. When the set size of memory items was below eight, the outer positions were filled with “X” as placeholders. The presentation of the memory items lasted for 2 seconds, allowing participants to read the letters and hear them spoken at the same time. A 3-second maintenance phase followed, where all the letters, but not the fixation point, disappeared. Afterwards, a probe was presented. The participants were instructed to remember the memory items and then indicate whether the probe belonged to the memory set or not with a button press. They were told to respond as quickly as possible without making errors. After the response, the participants received acoustic feedback on whether the response was correct. Participants performed two to eight sessions (4.5 sessions on average), each consisting of 50 trials.

#### Stereo-EEG Recording and Preprocessing

Between four and eight depth electrodes (each with eight 1.6 mm contacts spaced 5 mm apart) were stereotactically implanted in patients’ presumed epileptic area identified by the epileptologists. The electrode contacts were primarily located in the hippocampus and middle temporal gyrus (Fig. 3B). The data were recorded against a common intracranial reference with a sampling rate of 2000 Hz. For our analysis, we treated each hemisphere separately and excluded any hemisphere with fewer than three implanted electrodes. This criterion ensured that each included hemisphere contained between three and four electrodes. After the exclusion, we ended with 11 participants with left-hemispheric data and 9 for the right hemisphere. To utilize as many electrodes as possible, we did not remove the electrodes in the tissues that were later confirmed to be epileptogenic. A previous study using a subset of the dataset found that neurons within epileptogenic regions were also involved in working memory performance^49^.

The provided data were pre-segmented from 1s before the onset of encoding till 7s after. We down-sampled them to 1000 Hz and removed the trials of artifacts based on existing labels coming along with the dataset. No additional pre-processing was implemented.

### Traveling Wave Intensity Index

#### The relationship between TWII and the traveling wave pattern

We defined the region of interest (ROI) to be the space covered by nearby electrodes. In the present study, the ROIs were the temporal lobes in the left and right hemispheres. Assuming there is a group of neural waves, ***TW***, propagating through a limited area in the ROI (see Fig. 3C). The depth electrodes in the ROI are thus sampling ***TW*** along their orientations. We denote the traveling waves captured by individual depth electrodes as a vector ***TW_i_***. Its direction is the orientation of the electrode, and its magnitude is the consistency of phasic gradient on the electrode (see “*Computation of TWII and statistical testing for TWII existence*”, step 1 for computing ***TW_i_***). Fig. 3C shows three possible scenarios of ***TW_i_*** according to the electrode layout: (1) The most ideal case is when the electrode is completely positioned within the area with traveling wave and roughly parallel to the direction of ***TW***. The ***TW_i_*** would accurately capture ***TW***. (2) A less ideal case occurs when the electrode is only partially located in the traveling-wave area and / or its orientation moderately misaligned from direction of traveling waves. In this case, ***TW_i_*** would have a relatively small magnitude, and its direction would deviate moderately from the true traveling-wave direction. However, the deviation would never be large enough to reverse the direction of the estimated vector. (3) The most extreme case is when the electrode is perpendicular to the direction of traveling waves. The electrode would only capture apparent “standing waves” and yield ***TW_i_*** = 0, regardless of where the electrode is located relative to the traveling waves.

Since the second scenario is the most common case of depth electrode layout, taking the vectorial mean across all the ***TW_i_*** should increase the precision of estimation of ***TW***. Thus, we defined the Traveling Wave Intensity Index (TWII) by taking a power-normalized vectorial mean of all the TW_i_, as :

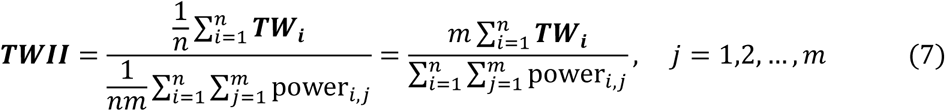

Here, power_i_ denotes the wave power measured at the *j*th contact of the *i*th electrode, and *m* is the number of contacts per electrode (here m = 8). The denominator represents the mean wave power across the ROI, thereby normalizing ***TWII*** by accounting for variability in wave amplitude.

The vector ***TWII*** is a value that can reflect the consistency of the traveling waves across all the electrodes in the ROI. Its direction approximates the direction of ***TW*** under the constraint of electrode orientation. The magnitude |*TWII*| has a range of 0, ∞, with values near zero indicating the absence of coherent traveling wave activity (including when there are only standing waves) and larger values indicating more consistent traveling wave patterns. This enables further statistical tests for the existence of traveling waves (see the section below).

A prerequisite of computing ***TWII*** is a minimum of three non-parallel electrodes in each ROI to ensure adequate three-dimensional spatial sampling. In the current study, the data from three participants did not qualify for this prerequisite and were therefore excluded from the analysis. Notably, ***TWII*** captures only the dominant and spatially coherent planal wave component across the ROI. In cases where the region contains circular traveling waves or multiple localized planal traveling waves with distinct propagation directions, ***TWII*** may not reflect this heterogeneity. Thus, the spatial scale of TWII is inherently defined by the size of the ROI.

#### Computation of TWII and statistical testing for TWII existence

Extended Data Fig. 1 depicts the full procedure of computing TWII, as described as follows:

*STEP 1: Quantify* **TW_i_**. The projected traveling waves component **TW_i_** on the *i*th electrode was computed based on a previously established method ^13,21^. The time series data recorded from each electrode contact was arranged according to their spatial layout. This resulted in a two-dimensional array of space and time. If there were oscillations in the ROI, one would see a strip-like pattern in the data. Depending on the waves’ propagation speed between the contacts, the strips would tilt in different directions. If the waves travel slowly, revealing consistent phasic shifts between the contacts, one would see tilted strip patterns. We used the Two-Dimensional Fast Fourier Transform (2DFFT) to detect these patterns and quantify temporal frequency and the traveling profile of the waves. To facilitate the computation, two processes were performed on the data: first, we linearly interpolated the data along the space dimension to nine contacts (originally eight). This is to ensure an odd number of data points in the spatial dimension so that we can separate the standing wave components in 2DFFT. Secondly, we applied a 500-ms sliding window moving through the time dimension of data in steps of 100 ms. By doing these, we segment data into multiple arrays of nine contacts × 500 ms and then applied 2DFFT on them. In the resulting power spectrum, the x-axis denotes the temporal frequency (how many cycles the waves reveal per second) and the y-axis denotes the spatial frequency (how many cycles the waves go through per contact). Because of the conjugation symmetry feature of 2DFFT, we consider the waves traveling along the electrode (from contact 1 to contact 8), *W*_+_, to be reflected in the top right and down left quarter, and the waves traveling against the electrode (from contact 8 to contact 1), *W*_–_, to be reflected in the top left and lower right quarter. The horizontal midline quantifies the power of standing waves on the electrodes. We restricted our analysis scope from 1 to 60 Hz. For each decomposed temporal frequency (hereafter refer to as frequency), we took the maximum value for each traveling direction, *W*_+,*max*_ and *W*_–,*max*_, in the corresponding column and quarter to represent their power. Hence, for each time and frequency point, we define its net traveling wave component projected on the electrode i as:

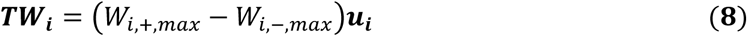

Here, ***u_i_*** denotes a unit vector estimated by applying singular value decomposition to the MNI coordinates of contacts on the electrode i. It points along the longitudinal axis of the electrode (from contacts 1 to 8). In other words, ***u_i_*** represents the direction of the electrode. These unit vectors were treated as free vectors, such that only their orientations, and not their spatial locations are considered in the following analysis.

*STEP 2: Compute **TWII**.* After acquiring the ***TW_i_*** from all the electrodes in the ROI (step 2 in Extended Data Fig. 1), one can compute the ***TWII*** with the formula (7). The ***TWII*** were then averaged across trials. Thus, there was one averaged ***TWII*** for each participant, time, and frequency point,

*STEP 3: Test whether there were consistent traveling wave patterns across participants.* For each time and frequency point, we then applied a null-model-based weighted 3D Raighley test on the averaged ***TWII*** across all participants. We developed it as a three-dimensional extension of the weighted Raighley test^32^. It is non-parametric and tests whether there is a consistent ***TWII*** direction across participants, weighted by the ranked norm of ***TWII***. To compute the statistics *R*^∗^, we first ranked the averaged ***TWII*** from each participant according to its norm |***TWII***|: the ***TWII*** with the smallest norm was given rank r = 1, the second smallest was given rank r = 2… and the largest was given the rank r = N (the total number of participants). Let

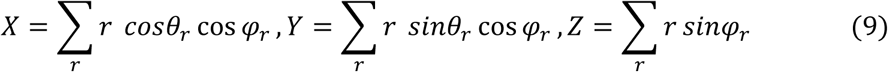

Where *θ* and *φ* are the azimuth and the elevation of the ***TWII***, respectively. Then we can define *R*^∗^ as

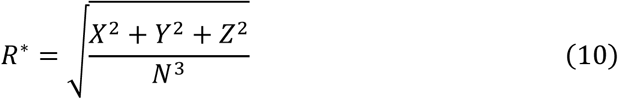

We obtained the null distribution of *R*^∗^ by simulating N (for left hemisphere N = 11, and right hemisphere N = 9) vectors randomly drawn from a spherical distribution 1,000,000 times. ***TWII*** that yield a *R*^∗^ larger than the top 5% of the simulated null *R*^∗^ distribution is considered to have a significantly consistent traveling wave direction across participants.

*STEP 4: Cluster-based correction for multiple comparisons*. Since we do not have an a priori estimate of the frequency and time points at which traveling waves would emerge, we computed TWII for the whole frequency spectrum and all time windows. As a result, each time-frequency point has its own ***TWII***, and the weighted 3D Raighley test was performed separately for each time-frequency point. To control for multiple-comparisons we followed the suggestion of Maris and Oostenveld^50^ and performed cluster-based correction. Importantly, the weighted 3D Raighley is a one-sample test and the input data does not come with permutable labels (e.g., experimental condition) to construct a null distribution like the examples given by Maris and Oostenveld^50^. To address this, we created the null distribution by permuting the spatial order of the contacts within the electrodes and repeating steps 1 to 3 for 1000 times. The rest of the correction followed what was recommended in Maris and Oostenveld^50^; We computed the sum of statistics *R*^∗^ for every significant time-frequency cluster for permuted and real data. Clusters with summed *R*^∗^ higher than the top 5% of its null distribution derived from permuted data were considered significant.

## Data and Code availability

The dataset reanalyzed by the current study is from Dimakopoulos et al.^30^, accessible at https://openneuro.org/datasets/ds004752/versions/1.0.1.

All data were analyzed with custom scripts in MATLAB R2023a (The MathWorks, Natick, MA, USA), and MATLAB Toolboxes, including: Image Processing Toolbox, Parallel Computing Toolbox, Signal Processing Toolbox, and Statistics and Machine Learning Toolbox. Scripts for artificial neural network model are available at https://github.com/artipago/traveling-waves-shields-pattern-encoding-/tree/publish. Scripts for computing TWII are available at https://anonymous.4open.science/r/TWII-2F76/README.md.

## Declaration of LLM usage

ChatGPT (OpenAI; GPT-4) were used to assist with generating code for supporting aspects of this work, including data simulation for TWII and data visualization, and for copy-editing the manuscript to improve language and readability. All core data analyses and statistical procedures, as well as the scientific interpretation of the results, were developed and conducted by the authors without the use of LLMs. All LLM-generated or edited content was reviewed and verified by the authors.

## Acknowledgement

We thank Michael Emil Aristodemou for the support in statistical consulting and Larrisa Behnke for suggestions on manuscript editing. We also thank Tzvetan Popov, Nicolas Langer, and Mike X Cohen for valuable insight and feedback on the traveling waves computation.

## Author Contribution

Y.Z. contributed to Conceptualization, Formal Analysis, Methodology, Software, Visualization, Writing – original draft, and Writing – review & editing. Y.H. contributed to Conceptualization, Formal Analysis, and Writing – review & editing. C.P. contributed to Conceptualization, Writing – original draft, and Writing – review & editing. P.T. contributed to Conceptualization and Writing – review & editing. L.I. contributed to Data Curation, Investigation, and Writing – review & editing. J.S. contributed to Conceptualization, Data Curation, Investigation, Methodology, and Writing – review & editing. A.A. contributed to Conceptualization, Formal Analysis, Funding Acquisition, and Writing – review & editing. P.S. contributed to Conceptualization, Funding Acquisition, Supervision, Writing – original draft, and Writing – review & editing.

