## Supplementary Materials for "Traveling brain waves support flexible storage of human working memory"

### Supplementary Analysis

**Speed of traveling waves along electrodes**

One theoretical possibility is that the observed traveling waves were artefacts introduced by electrode insertion, creating a conductive tunnel facilitating the propagation of electrical signals. To rule this out, we estimate the speed at which the waves traveled along individual electrodes. If the traveling waves were an artefact, one would expect a volume-conduction-like traveling pattern at the speed of light in brain tissue ($\approx 2 \times 10^8$ m/s). However, this was not the case (Extended Data Fig 2B). On average, in the left hemisphere the theta waves traveled at 0.18±0.10 m/s (M±SD), the left gamma waves traveled at 1.09±0.58 m/s, and the beta waves in the right hemisphere traveled at 0.41±0.26 m/s. Overall, the waves traveled at a speed many orders of magnitude below the speed of light, and aligned with the traveling waves' speed typically observed at the mesoscopic level (0.1-0.8 m/s[1]).

The estimation of the wave speed is as follows. During the first step of TWII computation, the traveling wave intensity along individual electrodes $W_{+}$ and $W_{-}$was defined as the maximum values for each temporal frequency column in the 2DFFT spectrum (Extended Data Fig 1, ①). We then took the spatial frequency index i (i = 1, 2, 3, 4) corresponding to this maximum value (If $W_{+} > W_{-}$, we took the spatial frequency index of $W_{+}$, and vice versa) to compute the speed, as

$$\mu = \frac{(n-1)Df}{i}$$

Where n is the number of interpolated contacts (here n = 9), D is the distance between contacts (here D = 0.005m), and f is the temporal frequency of oscillations. Because of the discrete nature of 2DFFT and the sparse spatial sampling of the contacts, the resulting speed estimation formed a discrete distribution with a range of $[2fD, 8fD]$, where its lower bound is limited by spatial Nyquist ($k_{max} = \pi/D$) and the upper bound is limited by the number of contacts .

**Regional oscillatory power confirms the presence of theta and beta waves**

The robustness of phasic analysis of oscillatory signals, including traveling wave computation, relies on the strength of oscillatory power. To verify the presence of oscillatory signals in the dataset, we calculated the regional oscillatory power. This is done by averaging the power spectra obtained via one-dimensional Fast Fourier Transform across all recording contacts in each hemisphere. We took the resulting power spectrum to fit the 1/f aperiodic components with the FOOOF toolbox (https://fooof-tools.github.io/fooof/). Results are illustrated in Extend Data Fig 2A. For the left hemisphere, two peaks of oscillatory power were identified as exceeding the aperiodic fit (Peak 1, Center Frequency [CF] = 8.90, Power [PW] = 0.17, Bandwidth [BW] = 5.74; Peak 2, CF = 18.44, PW = 0.13, BW = 12; Overall goodness of fit $R^2$ = 0.999; Error = 0.02). In more detail, the first peak corresponds to the theta (4-10 Hz) intracranial traveling waves observed in the left hemisphere (Fig. 4A). We did not find a peak in power in the frequency band corresponding to the gamma intracranial traveling waves (32-54 Hz). However, this is more likely to be due to underfitting of aperiodic components in the high frequency range (note the deviation of the aperiodic fit and original spectrum) than the absence of gamma oscillatory signals. In the right hemisphere, one oscillatory peak was identified (Peak 1, CF = 13.28, PW = 0.17, BW = 12.00; Overall goodness of fit $R^2$ = 0.998; Error = 0.02), aligned with the beta intracranial traveling waves (14-26 Hz) spotted in the right hemisphere (Fig. 4A).

Our analysis of the oscillatory power dynamic across the trial also showed that the traveling waves pattern is independent from the variation of the waves' oscillatory power. In Fig 4C we present the z-scored oscillatory power, computed as

$$Z_{t,f} = \frac{P_{t,f} - \mu_f}{\sigma_f}$$

where $P_{t,f}$ is the oscillatory power of frequency f at time t, and $\mu_f$ and $\sigma_f$ are the mean and standard deviation of the power across all time points. Our descriptive results show that the z-scored oscillatory power increased across nearly all frequency bands during encoding and retrieval, with a pronounced ramping at the retrieval onset and the encoding offset (note that each time point represents the starting point of a sliding window of 500 ms). During the maintenance phase, oscillatory power was consistently suppressed across the entire spectrum. Such patterns of oscillatory power differ from the sustained theta and gamma traveling waves or the semi-bursting beta traveling activities during maintenance and retrieval phases (Fig 4A).

### Extended Data Figure Legend

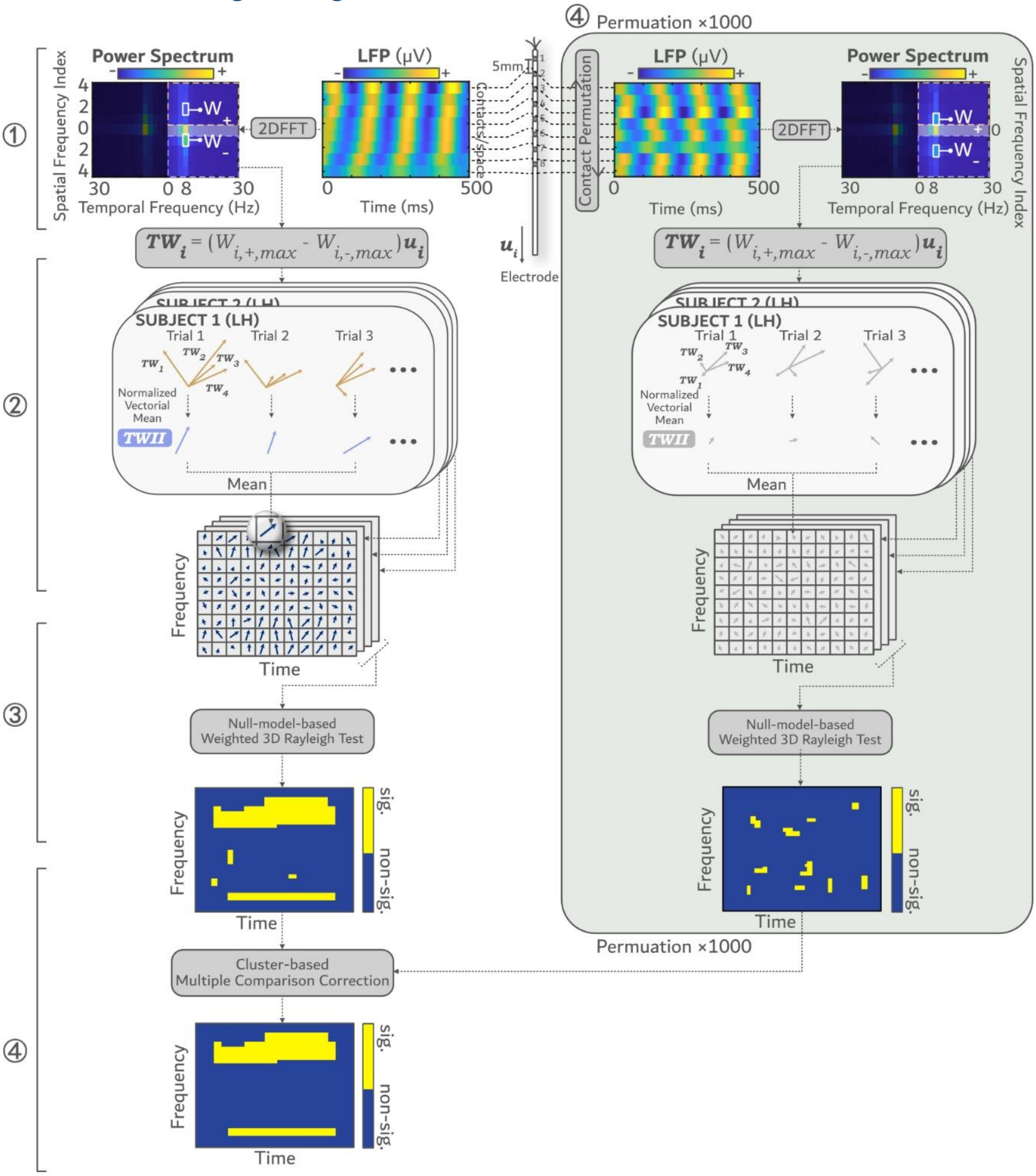


Extended Data Figure 1. The procedure of computing the Traveling Wave Intensity Index (TWII). More details see method.

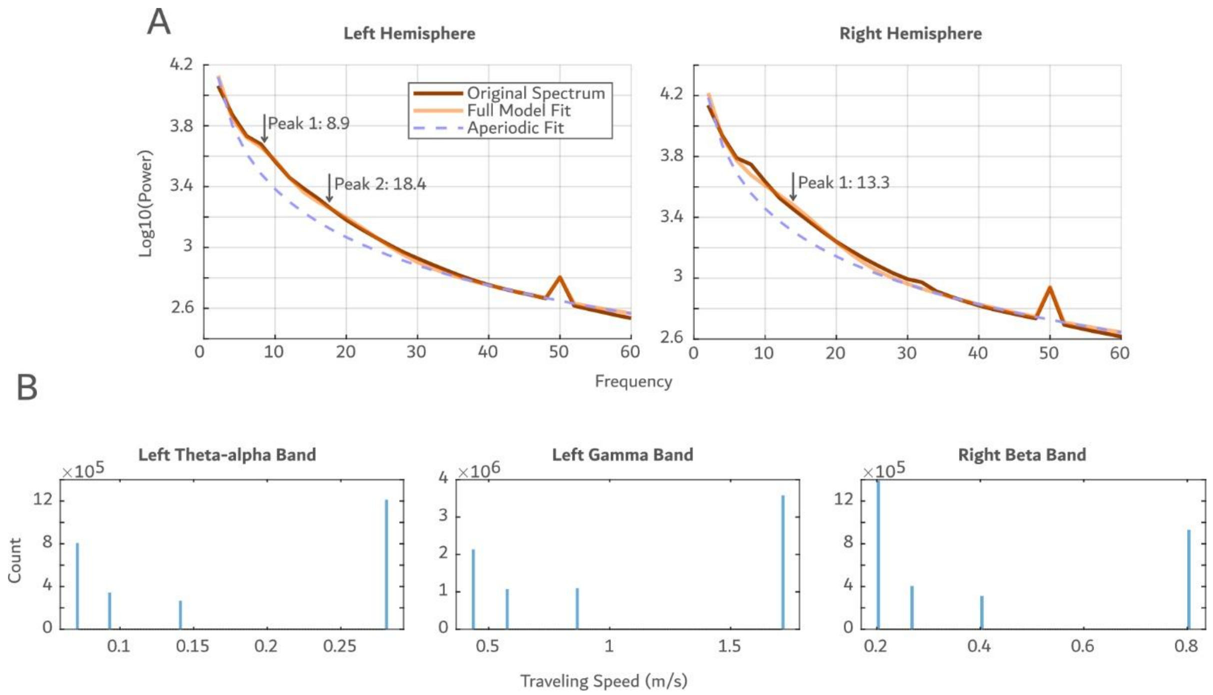


Extended Data Figure 2. The descriptive results of oscillatory power. (A) The power spectrum, superimposed on the 1/f aperiodic component and the full model fit provided by the FOOOF toolbox. The arrow indicates the peaks in the power spectrum and their corresponding frequencies that rise above the aperiodic line. (B) The distribution of traveling speeds for theta waves (left), gamma waves (middle) in the left hemisphere, and beta waves in the right hemisphere (right).
